# Suppression of cortico-striatal circuit activity improves cognitive flexibility and prevents body weight loss in activity-based anorexia in rats

**DOI:** 10.1101/2020.03.29.015115

**Authors:** Laura K Milton, Paul N Mirabella, Erika Greaves, David C Spanswick, Maarten van den Buuse, Brian J Oldfield, Claire J Foldi

## Abstract

**Background:** The ability to adapt behavior to changing environmental circumstances, or *cognitive flexibility*, is impaired in multiple psychiatric conditions, including anorexia nervosa (AN). Exaggerated prefrontal cortical activity likely underpins the inflexible thinking and rigid behaviors exhibited by patients with AN. A better understanding of the neural basis of cognitive flexibility is necessary to enable treatment approaches that may target impaired executive control.

**Methods:** Utilizing the activity-based anorexia (ABA) rat model and touchscreen operant learning paradigms, we investigated the neurobiological link between pathological weight loss and cognitive flexibility. We used pathway-specific chemogenetics to selectively modulate activity in neurons of the medial prefrontal cortex (mPFC) projecting to the nucleus accumbens shell (AcbSh) in female Sprague-Dawley rats.

**Results:** DREADD-based inhibition of the mPFC-AcbSh pathway prevented weight loss in ABA and improved flexibility during early reversal learning by reducing perseverative responding. Modulation of activity within the mPFC-AcbSh pathway had no effect on running, locomotor activity or feeding under ad libitum conditions, indicating the specific involvement of this circuit in conditions of dysregulated reward.

**Conclusions:** Both attenuation of weight loss in ABA and improved cognitive flexibility following suppression of mPFC-AcbSh activity aligns with the relationship between disrupted prefrontal function and cognitive rigidity in AN patients. The identification of a neurobiological correlate between cognitive flexibility and pathological weight loss provides a unique insight into the executive control of feeding behavior. It also highlights the utility of the ABA model for understanding the biological bases of cognitive deficits in AN and provides context for new treatment strategies.

## Introduction

The ability to modify cognitive strategies in novel or uncertain contexts in order to adapt to changing environmental demands is a necessary behavior exhibited by humans and animals. This *cognitive flexibility* is impaired in multiple psychiatric conditions, including anorexia nervosa (AN) and obsessive-compulsive disorder (OCD) [1]. These disorders commonly co-occur [2, 3], have highly concordant genetic etiologies [4, 5] and it has been suggested that OCD is a risk factor for the development of AN [6]. Engagement in rigid, rule-bound behavior, avoidance of novel situations and perseverative thinking are the behavioral hallmarks of cognitive inflexibility in both disorders. In patients with AN these manifest specifically as a preoccupation with food and body shape as well as excessive exercise to further the relentless pursuit of weight loss [3, 7, 8]. These observations reflect strict cognitive control over behavior which may be driven by heightened activity in the prefrontal cortex, as demonstrated by functional imaging studies in patients with AN when participating in reward-related tasks [9]. Clinically, cognitive flexibility is assessed through tests of executive function including attentional set-shifting, response inhibition and reversal of reward-based associations. Impairments in the ability to adapt behavior in these tasks in response to changing rules have been consistently demonstrated in patients with AN, who make more perseverative errors compared to healthy controls [10–12]. However, because starvation itself leads to food obsessions and increased cognitive rigidity independent of clinical AN [13], it is important to differentiate between alterations that precede the onset of the disorder (and thus contribute to increased risk) and those that are secondary to starvation (that may play a role in reinforcing the behavioral manifestations of AN). In support of cognitive rigidity as an etiological risk factor for AN, impaired set-shifting has been demonstrated in healthy sisters of AN patients [14–16], persists after body weight recovery [16, 17] and is independent of body weight status and duration of illness [15, 18].

Interrogation of the neurobiological underpinnings of AN, particularly those that precede onset of the disorder, is difficult in humans. To this end, the activity-based anorexia (ABA) rodent model has been utilized effectively over several decades [19] and remains the only experimental approach whereby laboratory animals will choose self-starvation over homeostatic balance [20]. In the ABA paradigm, when time-limited access to food is paired with access to a running wheel, rats (and mice) reduce food intake and dramatically increase running activity, producing precipitous and rapid body weight loss that will lead to death if unchecked [21]. Voluntary access to a wheel is critical to the development of the ABA phenotype, without which animals quickly learn to increase intake when food is available to maintain body weight [22]. Hyperactivity in response to restricted food availability, while extreme in ABA, mimics, in its initial stages, the adaptive response of rodents in the wild to periods of food scarcity [23]. Here, strategies to maintain homeostatic energy balance and survive have to consider both reducing activity levels to compensate for a reduced energy intake and increasing activity levels to effectively forage [23–25]. While increasing activity in order to find food may be appropriate for rodents in the wild, it is maladaptive in a laboratory setting – this is certainly the case in ABA.

Hyperactivity in response to food scarcity (known as “starvation-induced hyperactivity”) is often considered a compulsive behavior [26] and occurs in the majority of animals exposed to ABA conditions. Notably, recent evidence suggests that not all animals are susceptible to body weight loss in ABA and in fact there exists a distinct population of rats [27] and mice [28] that are resistant to developing the hyperactive phenotype. The propensity for excessive wheel running in animals susceptible to ABA is thought to be associated with an inability to flexibly adapt to changes in feeding schedules [29]. The question then becomes whether this apparent breakdown of cognitive-behavioral flexibility is a cause or a consequence of the tendency to run rather than eat in the ABA paradigm. Indeed, ABA is shown to produce deficits in reversal learning in rats compared to their control counterparts with limited food access but without access to a running wheel [26]. This effect was no longer observed following weight recovery, suggesting, in this instance, it is a consequence rather than a driving factor in susceptibility in pathological body weight loss in ABA. However, reversal learning was not impaired in body weight-matched controls, suggesting the inflexible phenotype cannot be explained by the acute effects of starvation in this study. Therefore, inflexibility must be related, at least in part, to the development of compulsive wheel running that accompanies exposure to ABA conditions. However, what is currently unknown is the extent to which cognitive rigidity is involved in the *generation* of the ABA phenotype.

Tests of cognitive flexibility in rats involve extended training using food rewards to shape behavior. This is problematic for the investigation of the cognitive antecedents to ABA not only because of the central role of reward processes in the development of ABA [20, 30] but also because repeated handling may itself influence susceptibility to body weight loss in the paradigm [31]. One solution is to investigate the neural correlates of cognitive-behavioral flexibility in the generation of the ABA phenotype. Executive functions, including cognitive flexibility, are broadly under the control of the prefrontal cortex that has widespread connectivity with subcortical regions [32] and contributes to feeding and food-seeking behavior [33, 34]. Of particular interest to AN and ABA is the interaction between the medial prefrontal cortex (mPFC) and the ventral striatum, which is critical for the higher-order (top-down) control of reward-based feeding [35, 36] as well as components of cognitive-behavioral flexibility [37, 38]. In fact, impaired cognitive flexibility in patients with AN seems to be directly associated with altered activity of fronto-striatal networks [39]. The nucleus accumbens shell (AcbSh) is a sub-region of the ventral striatum that integrates reward and motivational inputs relevant to feeding behavior [40] and activation of the mPFC suppresses AcbSh-evoked feeding behavior [41]. Signaling in the AcbSh also contributes to the development of ABA in rats, whereby chemogenetic activation of ventral tegmental area (VTA) neurons with direct projections to the AcbSh can both prevent and reverse the ABA phenotype [42]. Whether or not the prefrontal inputs to ventral striatal pathways are directly involved in pathological body weight loss remains to be determined.

In the present study, we sought to ascertain a) if chemogenetic modulation of the mPFC-AcbSh projection pathway impacts on body weight, feeding behavior and wheel running activity in ABA and b) whether this same neuronal projection is involved in cognitive-behavioral flexibility.

## Materials and Methods

### Pathway-specific chemogenetics

A retrogradely-transported adeno-associated virus carrying Cre recombinase (AAV pmSyn1-EBPF-Cre, abbreviated retro-Cre; Addgene viral prep #51507-AAVrg; a gift from Hongkui Zeng) was injected into the medial shell of the nucleus accumbens (AcbSh). In order to specifically target the mPFC to AcbSh pathway, Cre-dependent DREADD constructs were injected into the mPFC, a well-defined source of afferent input to the AcbSh. These were either excitatory [pAAV-hSyn-DIO-hM3D(Gq)-mCherry, abbreviated hM3Dq; Addgene viral prep #50474-AAV5], inhibitory [pAAV-hSyn-DIO-hM4D(Gi)-mCherry, abbreviated hM4Di; UNC Vector Core, AAV5] or contained only the mCherry fluorophore as a control (pAAV-hSyn-DIO-mCherry, abbreviated mCherry; Addgene viral prep #50459-AAV5; a gift from Bryan Roth). The activity of the mPFC-AcbSh pathway was then modulated using the synthetic ligand clozapine N-oxide (CNO; Carbosynth Ltd, Berkshire UK). CNO was dissolved in 5 % DMSO, diluted in 0.9 % saline and administered intraperitoneally to all animals (hM3Dq; 0.3 mg/kg, hM4Di; 3.0 mg/kg. mCherry; both doses, counterbalanced) 30 min prior to the onset of the feeding period or behavioral intervention and 90 min prior to transcardial perfusion.

### Animals and housing

Female Sprague-Dawley rats (Animal Resources Centre; WA, Australia) were used in all experiments. The number of animals per groups and body weight ranges are indicated in **Table 1**. All experimental procedures on animals were approved by the relevant Monash (ERM 15171) or La Trobe (AEC18-17) Universities Animal Ethics committees.

**Table 1:**
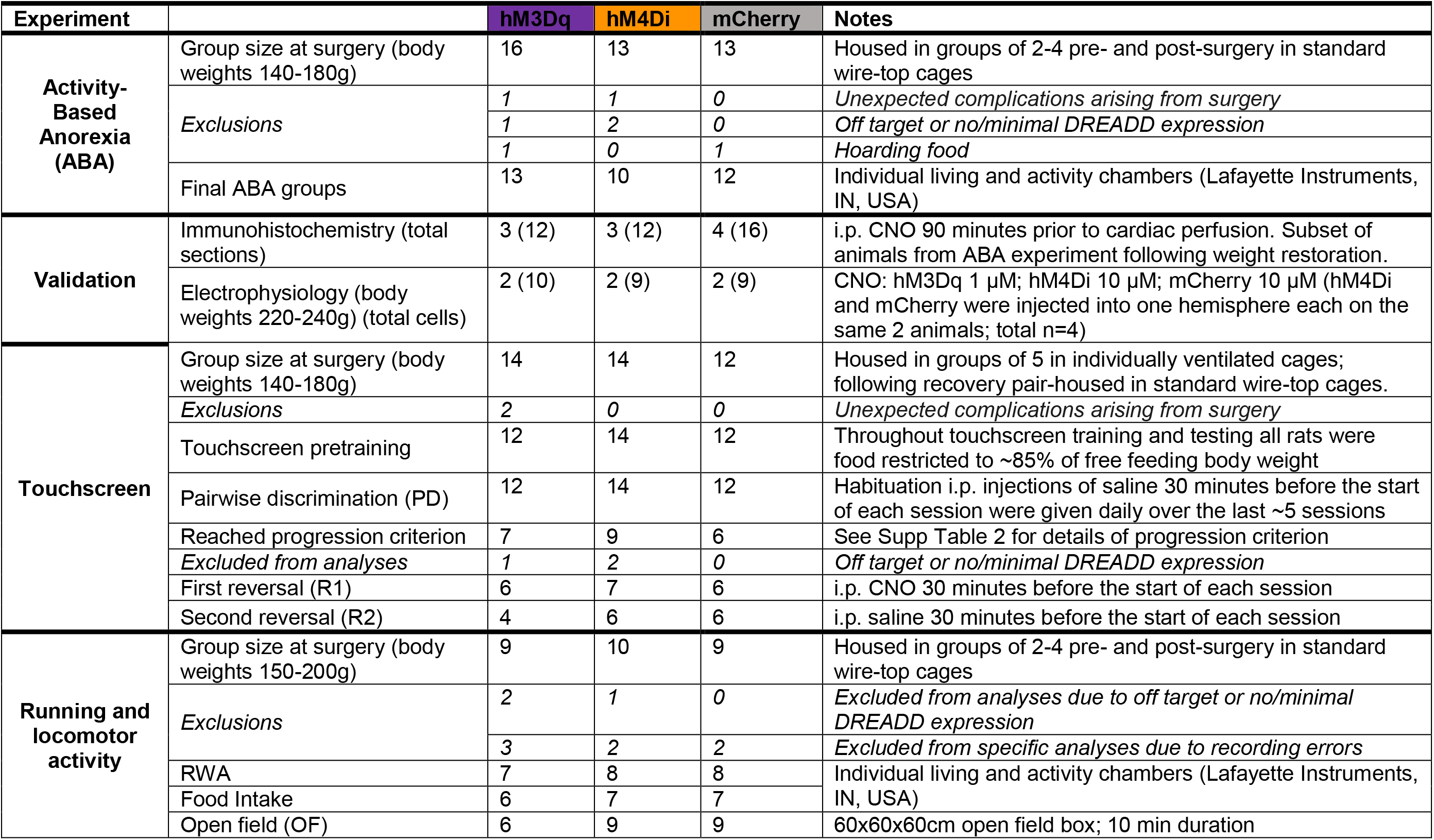
Animals and housing for each experiment

### Surgical and viral injection procedures

All rats were injected bilaterally with 200 nl of retro-Cre (6 × 10^12^ genomic copies/μl) into AcbSh (from bregma: AP: +1.6 mm, ML: ± 0.8 mm, DV: −6.6 mm) using a stereotaxic apparatus (Kopf Instruments, CA, USA). Rats then received bilateral injections of 200 nl of DREADD viral preparation (7 x 10^12^ genomic copies/μl) into the mPFC (from bregma; AP: +2.9 mm, ML: ± 0.5 mm, DV: −3.3 mm). Viruses were injected through pulled borosilicate glass micropipettes (Drummond Scientific; PA, USA) with tip diameters ~40 μm. Both viruses were injected under the same anesthetic (2.5 % isoflurane in oxygen) over 5 min, with pipette tips left at coordinates for an additional 5 min after each infusion to allow virus dispersal. Meloxicam (2 mg/kg, s.c.; Boehringer Ingelheim, Germany) was administered during surgery and Enrofloxacin (0.05 mg/ml; Bayer, Germany) was delivered in the drinking water to all rats for 48 h post-surgery. Viral infusions were conducted at least 10 days prior to the experimental period, to allow for virus transport and recombination.

### Electrophysiological validation of DREADD receptor activity

Whole-cell current clamp recordings from mCherry-expressing mPFC neurons following CNO application were performed as described in **Supplementary Methods 1 & 2**.

### Activity-based anorexia (ABA)

The ABA phenotype was generated as previously described [42] except that animals were not allocated to treatment groups matched for baseline running activity because the groups were necessarily specified by the viral construct infused. Therefore, running wheel activity was analyzed for individual rats as “change from baseline” values to account for any group bias. The ABA protocol was maintained until rats reached < 80 % of baseline body weight or for a maximum of 10 days.

### Running and locomotor activity under *ad libitum* feeding conditions

To determine whether effects of mPFC-AcbSh circuit modulation on ABA-associated behaviors were specific to the paradigm, we assessed feeding, running wheel activity and general locomotor activity in a separate cohort of rats under *ad libitum* fed conditions. These animals underwent the same wheel habituation and CNO administration protocols as those exposed to ABA, followed by a 10 min trial in the open field. The open field was constructed of grey acrylic (60 x 60 x 50 cm) and tests were conducted in the light phase under normal (~80 lx) lighting conditions. The apparatus was cleaned with 10 % ethanol between trials, which were recorded with an overhead camera and tracked using EthoVision XT software (Noldus; Wageningen, The Netherlands).

### Discrimination learning and cognitive flexibility

A reversal learning task was used to assess cognitive flexibility in touchscreen operant testing chambers (see **Supplementary Methods 3**). The reversal learning task had three phases that were conducted over daily sessions (6 days/week) using 45 mg sucrose pellet rewards (Able Scientific; WA, Australia) until rats were able to complete 100 trials in less than 60 min with ≥ 80 % accuracy (i.e. progression criterion, see **Supplementary Tables 1 & 2**). To advance to the next phase each rat was required to reach the progression criterion, if they failed to do so after 30 sessions, or performed no better than chance after 20 sessions, they were removed from the experiment (**Supplementary Fig 3**).

### Immunohistochemistry and imaging

Following completion of behavioral assays, rats were euthanized with sodium pentobarbitone (Lethabarb; 150 mg/kg) and perfused transcardially with 200 ml 0.9 % saline followed by 200 ml 4% paraformaldehyde in phosphate buffer (PFA-PB). Brains were excised for immunohistochemical confirmation of the extent and location of DREADD expression. Following transcardial perfusion, brains were post-fixed in 4 % PFA-PB solution overnight at 4 °C, followed by submersion in 30 % sucrose in PB solution for 3-4 days. Brains were sectioned at 35 μm using a cryostat (CM1860, Leica Biosystems) for standard immunohistochemical processing using the primary antibodies mouse anti-DsRed (sc-390909, 1:1000, Santa Cruz, TX) and rabbit anti-cfos (sc-52, 1:2000, Santa Cruz, TX) overnight at room temperature. Subsequently, sections were incubated for 90 min in secondary antibodies conjugated with Alexa Fluor dyes 594 and 647 (1:1000, Abcam, UK). Imaging was conducted using a Leica SP5 confocal microscope (20X objective magnification, 1024 x 1024 resolution) running LAS AF software (Leica MicroSystems; Germany). Tiled, z-stacked images (4 x 2 tiles, *z*=9) were captured of the mPFC, from 4 x 35 μm sections from each animal and confirmation of the action of CNO was assessed by colocalization of mCherry with Fos using Image J software (National Institutes of Health, USA).

### Statistical analyses

Statistical analyses were performed with GraphPad Prism 8.0 (GraphPad Software; CA, USA) and significance for all tests was set at *p*<0.05. One-way ANOVA followed by Dunnett’s post-hoc multiple comparisons was used to analyze mCherry and Fos colocalization. Electrophysiological changes induced by CNO were assessed relative to baseline using paired t-tests. Survival in ABA was evaluated with a Log-rank (Mantel-Cox) χ^2^ test with Bonferroni’s correction for multiple comparisons. One-way ANOVAs were used to analyze mean daily ABA food intake, change in mean daily RWA and FAA, and open field distance moved. Mixed-effects models followed by Tukey’s post-hoc multiple comparisons were used to analyze touchscreen testing outcomes from the first 100 accuracy trials of each phase and number of trials to reach criterion. Fisher’s exact test was used to compare learning outcome of touchscreen images. Two-way RM ANOVA with Tukey’s post-hoc multiple comparisons were used to analyses ad libitum food intake and running wheel activity following CNO administration. Initial group sizes were based on power calculations prior to the experiments; some animals were excluded from the final analyses for reasons detailed in **Table 1**.

## Results

### Validation of CNO effects on DREADD-expressing mPFC-AcbSh projection neurons

In order to validate our viral infusion strategy (**Fig 1A**) and CNO-induced effects on behavioral outcomes, we first examined immunohistochemically the proportion of DREADD-expressing neurons that co-expressed cFos, a marker of recent neuronal activation (**Fig 1B**). Compared to mCherry-expressing controls, neurons expressing hM3Dq and hM4Di DREADDs showed increased (hM3Dq; *p*=.0139), and decreased (hM4Di; *p*<.0001) colocalization with cFos, respectively (**Fig 1C**). Using whole-cell patch clamp electrophysiology, we also demonstrated changes in membrane potential for hM3Dq- and hM4Di-expressing neurons upon application of CNO (hM3Dq; *p*<.0001, hM4Di; *p*=.0001), while CNO had no effect on the membrane potential of mCherry-expressing neurons (**Fig 1D**). Bath application of CNO increased the firing frequency of hM3Dq-expressing neurons (*p*=.0003) and the rheobase of hM4Di-expressing neurons (*p*<.0001), but had no effect on mCherry control cells (**Fig 1E-F**). Input resistance of hM3Dq-expressing neurons was increased following CNO application (*p*=.0066) but decreased in hM4Di-expressing cells (*p*=.0093), with no effect on mCherry-expressing cells (**Fig 1G**). Current-voltage relationships conducted for each group pre- and post-CNO application confirmed closure and activation of one or more K+ channels in hM3Dq- and hM4Di-expressing cells, respectively (**Supplementary Fig 1**). The CNO-induced excitation and inhibition were associated with a reversal potential of −96 ± 6.3mV (hM3Dq; *n* = 8) and −90 ± 9.9mV (hM4Di; *n* = 7) respectively, which approximates the reversal potential for potassium under our recording conditions. Representative whole-cell current clamp recordings of responses to CNO for each DREADD type are displayed in **Fig 1H**.

**Figure 1:**
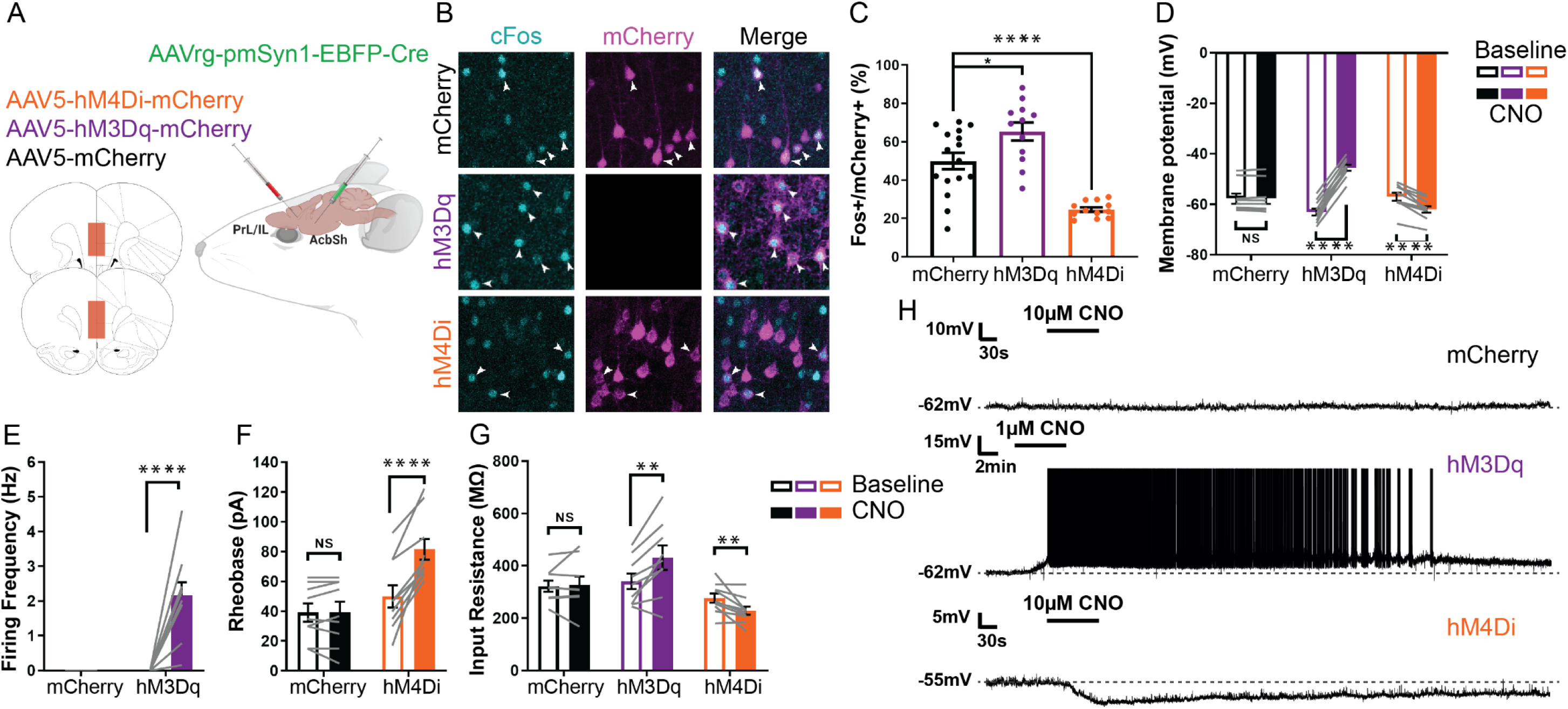
Validation of DREADD modulation of the mPFC-AcbSh projection neurons. CNO effects on DREADD-expressing neurons in CNO-treated animals **(A-C)** and after bath application of CNO **(D-H). A)** Schematic showing the dual viral injection strategy used to selectively target and manipulate neurons projecting from the mPFC (PrL/IL) to nucleus accumbens shell (AcbSh) with Cre-recombinase (green) mediated incorporation of mCherry (control; black), activating hM3Dq-mCherry DREADDs (purple), or inhibitory hM4Di-mCherry DREADDs (orange) into these neurons. Red boxes indicate where neuronal labelling was located 2.7 - 3.2 mm rostral from Bregma. **B)** Representative images of mPFC, collected 90 min after i.p. injection of CNO (0.3-3mg/kg). Fluorescent staining shows cFos (cyan), a marker of neuronal activation, and mCherry (magenta). Fos+ and mCherry+ colocalization is indicated with white arrow heads. **C)** Percentage of Fos+ and mCherry+ colocalization. One-way ANOVA: *F*(2, 37)=25.39, *p*<.0001. Post-hoc Dunnett’s multiple comparisons: hM3Dq > mCherry (*p*=.0139); hM4Di < mCherry (*p*<.0001). Bar charts representing the changes in **(D)** membrane potential (mV), **(E)** firing frequency (Hz), **(F)** rheobase (pA), and **(G)** input resistance (MΩ) when **(H)** CNO (1-10μM) was bath applied during *ex vivo* whole-cell patch clamp recordings of virally-transduced mPFC to AcbSh pyramidal neurons (mCherry, n=9; hM3Dq, n=10; hM4Di, n=10). Electrophysiology data were analysed with paired t-tests (within-groups). **D)** Membrane potential: mCherry *t(8)*=5.470, *p*>.9999; hM3Dq *t(9)*=20.65, *p*<.0001; hM4Di *t*(9)=6.525, *p*=.0001. **E)** Firing frequency: hM3Dq *t*(9)=5.669, *p*=.0003. **F)** Rheobase: mCherry *t*(8)=.06100, *p*=.9529; hM4Di *t*(9)=7.424, *p*<.0001. **G)** Input resistance: mCherry *t(8)*=0.3753, *p*=.7172; hM3Dq *t(8)*=3.644, *p*=.0066; hM4Di *t(9)*=3.295, *p*=.0093. Dots and lines represent individual values, bars are group means ± SEM. NS = not significant (*p*>.05), \**p*<.05, \*\**p*<.01, \*\*\**p*<.001, \*\*\*\**p*<.0001.

### Cortico-striatal circuit inhibition improves body weight maintenance in ABA

To evaluate the impact of mPFC-AcbSh modulation on body weight loss in ABA, rats were allowed 7 days to acclimate to the running wheel with ad libitum access to food (**Fig 2A1**) followed by a maximum of 10 days exposure to ABA conditions (**Fig 2A2**) with concomitant daily administration of CNO. Individual body weight loss trajectories were substantially attenuated following mPFC-AcbSh inhibition (**Fig 2B**), whereby all rats were able to maintain body weight above 80 % of baseline and were resistant to developing ABA, compared to rats that had this circuit activated (*p*=.0030) and control rats (*p*=.0243) (**Fig 2C**). An increase in food intake following mPFC-AcbSh inhibition (**Fig 2D**) was the main driver of improvements to body weight maintenance (*p*=.0252), with no corresponding increase in overall running activity (**Fig 2E**) or “food restriction-evoked hyperactivity” (i.e. the change in running activity elicited by restricted food access; **Fig 2F**) when this circuit was suppressed. However, mPFC-AcbSh inhibition specifically increased running in anticipation of food (food anticipatory activity; FAA **Fig 2G**, *p*=.0336). Conversely, stimulation of the mPFC-AcbSh exacerbated food restriction-evoked hyperactivity (**Fig 2F**, *p*=.0059), contributing to slightly poorer body weight maintenance throughout ABA exposure.

**Figure 2:**
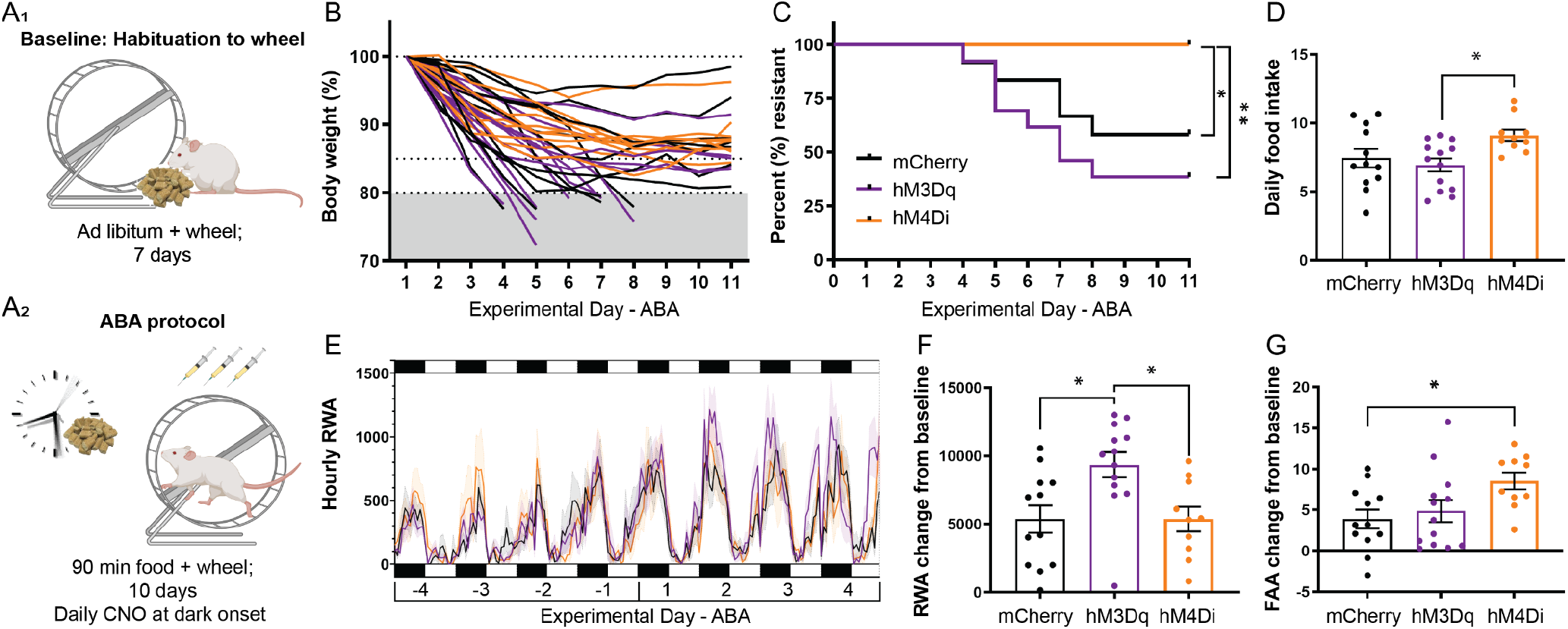
Effects of cortico-striatal circuit modulation on the development of ABA. Outcomes for rats exposed to the ABA protocol **(A)** expressing either only mCherry (black, n=12), activating hM3Dq-mCherry DREADDs (purple, n=12) or inhibiting hM4Di-mCherry-DREADDs (orange, n=10) in neurons projecting from the mPFC to the AcbSh. **B)** Individual body weight % loss trajectories; rats removed upon reaching <80%. **C)** Survival (%) was significantly greater for hM4Di than both mCherry (*p*=.0243) and hM3Dq (*p*=.0030); Log-rank (Mantel-Cox) χ^2^ tests using Bonferroni correction for multiple comparisons. **D)** Mean daily ABA food intake was significantly greater for hM4Di than hM3Dq (*p*=.0229) **E)** Hourly RWA over the last four days of habituation (−4 to −1) and first 4 days of ABA (1 to 4). Black boxes represent lights off. **F)** Increase in mean daily RWA from Habituation to ABA was significantly greater for hM3Dq than both mCherry (*p*=.0126) and hM4Di (*p*=.0180). **G)** Increase in mean daily food anticipatory activity (FAA) as a proportion of daily RWA from Habituation to ABA was significantly greater for hM4Di than mCherry (*p*=.0333). **D, F & G)** One-way ANOVA with post-hoc Tukey’s multiple comparisons, see **Supplementary Table 3** for details. **D-G)** Dots are individual animals, bars/lines are group means ± SEM (shaded bands in **E**). \**p*<.05, \*\**p*<.01

### Cortico-striatal circuit modulation does not impact general locomotor or feeding behavior

To determine effects of mPFC-AcbSh modulation under *ad libitum* fed conditions, rats were exposed to the same experimental protocol as for assessment of ABA, but in this case food access was not restricted (**Fig 3A**). While running activity and food intake increased from baseline for all groups during the CNO administration period (consistent with continuing access to the wheel; see **Supplementary Fig 2**), neither stimulation nor suppression of mPFC-AcbSh pathway neurons significantly altered food intake (**Fig 3B**) running activity (**Fig 3C**), FAA (**Fig 3D**) or general locomotor activity in an open field (**Fig 3E**) compared to mCherry controls.

**Figure 3:**
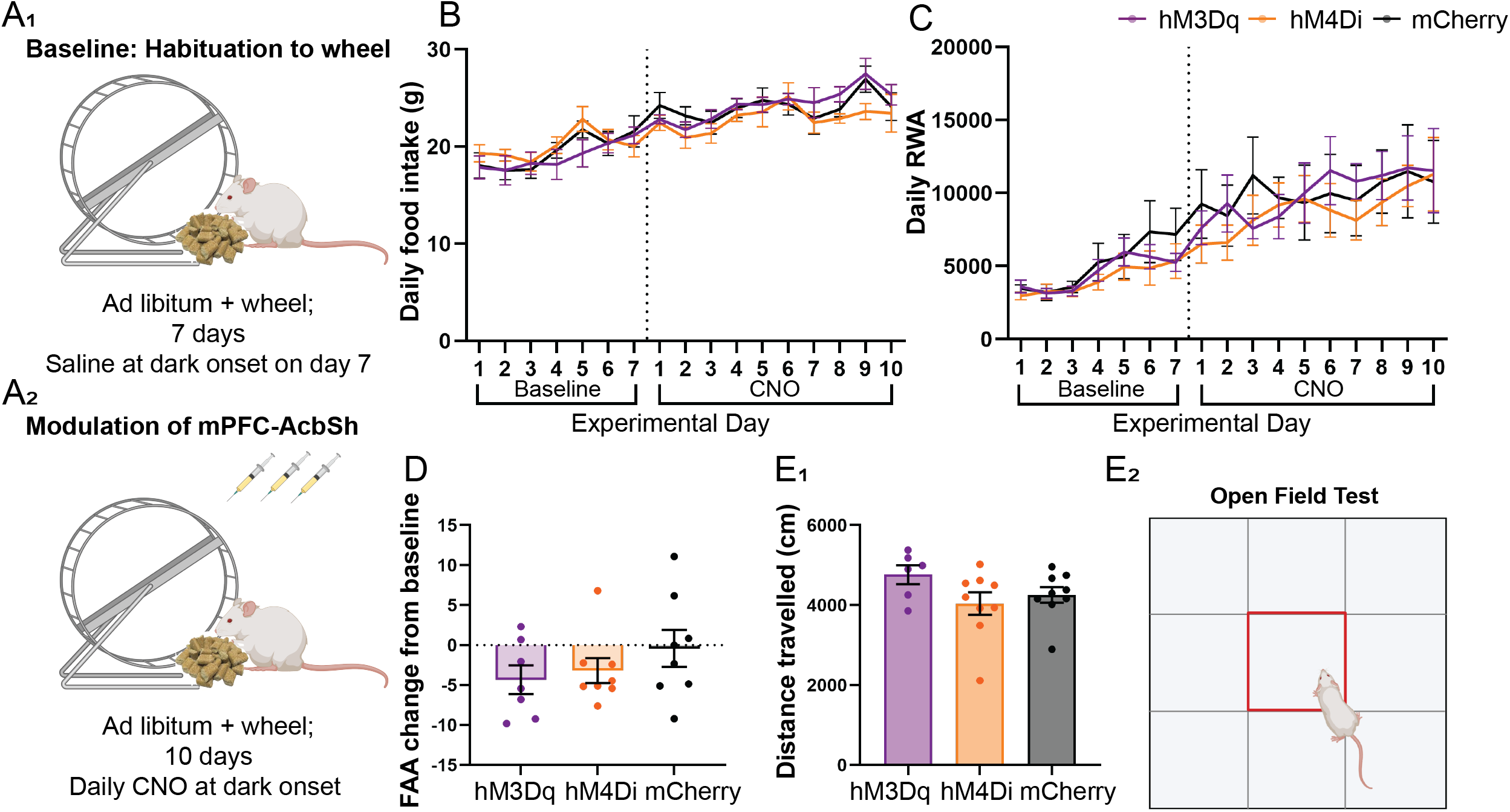
Effects of cortico-striatal modulation under ad libitum feeding conditions. Outcomes for rats maintained on ad libitum food access and exposed to wheel access and CNO administration that mirrors the ABA protocol **(A)** expressing either only mCherry (black, n=7-9), activating hM3Dq-mCherry DREADDs (purple, n=6-7) or inhibiting hM4Di-mCherry-DREADDs (orange, n=7-9) in neurons projecting from the mPFC to the AcbSh. Daily food intake **(B)** and daily running wheel activity **(C)** over 7 days of baseline and 10 days of CNO administration; mean ± SEM. **D)** Change in food anticipatory activity (FAA; RWA in the hour before lights off/food access in ABA as percentage of daily RWA) was not significantly different between groups (*F*[2, 20]=1.078, *p*=.3591). **E1)** Distance travelled (cm) in 10 minutes in the open field **E2)** was not significantly different between groups (*F*[2, 21]=1.920, *p*=.1715). **D and E1)** One-Way ANOVA; individual rats (dots, group mean (bars) ± SEM.

### Cortico-striatal circuit inhibition reduces perseverative responding in reversal task

Pairwise discrimination of visual stimuli and serial reversal of reward contingencies (**Fig 4A**) was utilized to examine the impact of modulation of mPFC-AcbSh activity on cognitive flexibility. There were no significant differences between groups in the ability of rats to learn the discrimination task or substantial changes induced by mPFC-AcbSh modulation to overall accuracy that determined progression through the reversal task (**Fig 4B**). However, in the early perseverative phase of visual reversal learning (first 100 reversal trials; see **Fig 4C2**), which coincided with initial activation or inhibition of the mPFC-AcbSh pathway, there was a significant improvement in accuracy following suppression of the circuit compared to both other groups (**Fig 4D**; hM4Di vs. mCherry *p*=.0366; vs. hM3Dq *p*=.0121). Activation of the mPFC-AcbSh pathway increased perseverative responding (**Fig 4E**, hM3Dq vs. mCherry *p*=.0346; vs. hM4Di *p*=.0025), demonstrated by a greater ratio of incorrect to correct responses (**Fig 4F**, hM3Dq vs. mCherry; *p*=.0382, vs. hM4Di; *p*=.0008). mPFC-AcbSh activation also increased response latency (**Fig 4H**, hM3Dq vs. mCherry; all *p*s≤.0193) and although animals in the hM3Dq group were slower to collect sucrose rewards (*p*=.0434), this was not specifically associated with activation of the circuit (**Fig 4l**). Similarly, animals with hM3Dq activation required fewer trials to reach criterion in both first and second reversals, indicating an unintended group bias rather than an effect mediated by mPFC-AcbSh stimulation (correct trials *p*s≤.0350, total trials *p*s≤.0468; **Figure 5**).

**Figure 4:**
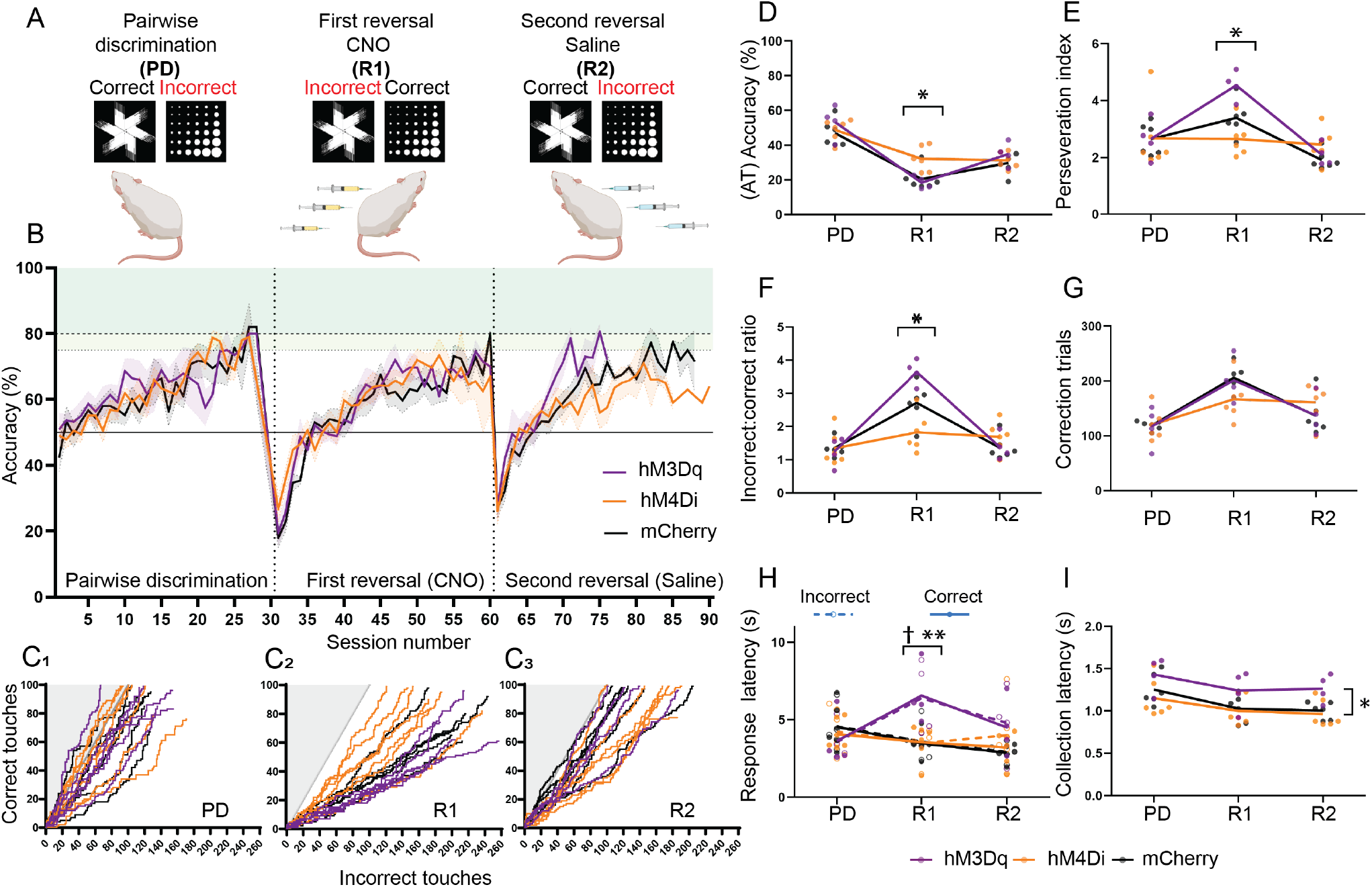
Effects of cortico-striatal circuit modulation on touchscreen visual discrimination and reversal learning. Touchscreen reversal learning performance measures for rats expressing either only mCherry (black, n=6), the activating hM3Dq DREADD (purple, n=6) or the inhibiting hM4Di DREADD (orange, n=7) in neurons projecting from the mPFC to the AcbSh. Schematic overview of (**A**) and accuracy (% correct in accuracy trials) over (**B;** group means ± SEM [shading]) the three phases of the reversal learning protocol (Pairwise Discrimination PD; First Reversal [R1]; Second Reversal [R2]). **C)** Individual rats’ progressive performance across the first 100 accuracy trials in each phase. Incorrect touch → X+1; Correct touch → Y+1 (max 100). Grey line shows chance performance with incorrect:correct ratio of 1.0. **D-I)** Individual animals (dots) and group means (lines) for the first 100 accuracy trial block in each phase. Only rats that progress to R2 are included (mCherry n=6, hM3Dq n=4, hM4Di n=6). Mixed-effect analysis followed by Tukey’s post-hoc multiple comparisons, see **Supplementary Table 3** for details. **D)** Accuracy % was significantly different between phases (*p*<.0001). Accuracy in R1 was significantly greater for hM4Di than both mCherry (*p*=.0366) and hM3Dq (*p*=.0121). **E)** The average number of consecutive incorrect responses (perseveration index [PI]) was significantly different across phases (*p*<.0001), with a significant phase*group interaction (*p*=.0048). PI in R1 was significantly greater for hM3Dq than both hM4Di (*p*=.0025) and mCherry (*p*=.0346). **F)** The ratio of incorrect to correct touches was significantly different between phases (*p*<.0001), with a significant phase*group interaction (*p*<.0001). Incorrect:correct ratio at R1 was significantly higher for hM3Dq than both hM4Di (*p*=.0008) and mCherry (*p*=.0382). **G)** The number of correction trials was significantly different across phases (*p*<.0001). **H)** There were significant phase*group interactions in the latency to touch the correct image (solid circles and lines, *p*=.0034) and the incorrect image (unfilled circles and dashed lines, *p*=.0051), with hM3Dq rats taking significantly longer than both hM4Di and mCherry rats to make both correct (asterisks; *p*=.0018 and *p*=.0014 respectively) and incorrect (cross *p*=.0148 and *p*=.0193 respectively) responses. **I)** The latency between touching the correct image and collecting the rewarded sucrose pellet (reward latency) was significantly different between phases (*p*<.0001). There was also a significant main effect of group (*p*=.0434; hM3Dq > hM4Di *p*=.0404), with hM3Dq reward latency significantly longer than hM4Di in R2 (*p*=.0365). Black vertical brackets show significant main effect group differences. Black horizontal brackets show significant between group comparisons within the enclosed phase (i.e. simple effects within phase). † and \**p*<.05, \*\**p*<.01.

**Figure 5:**
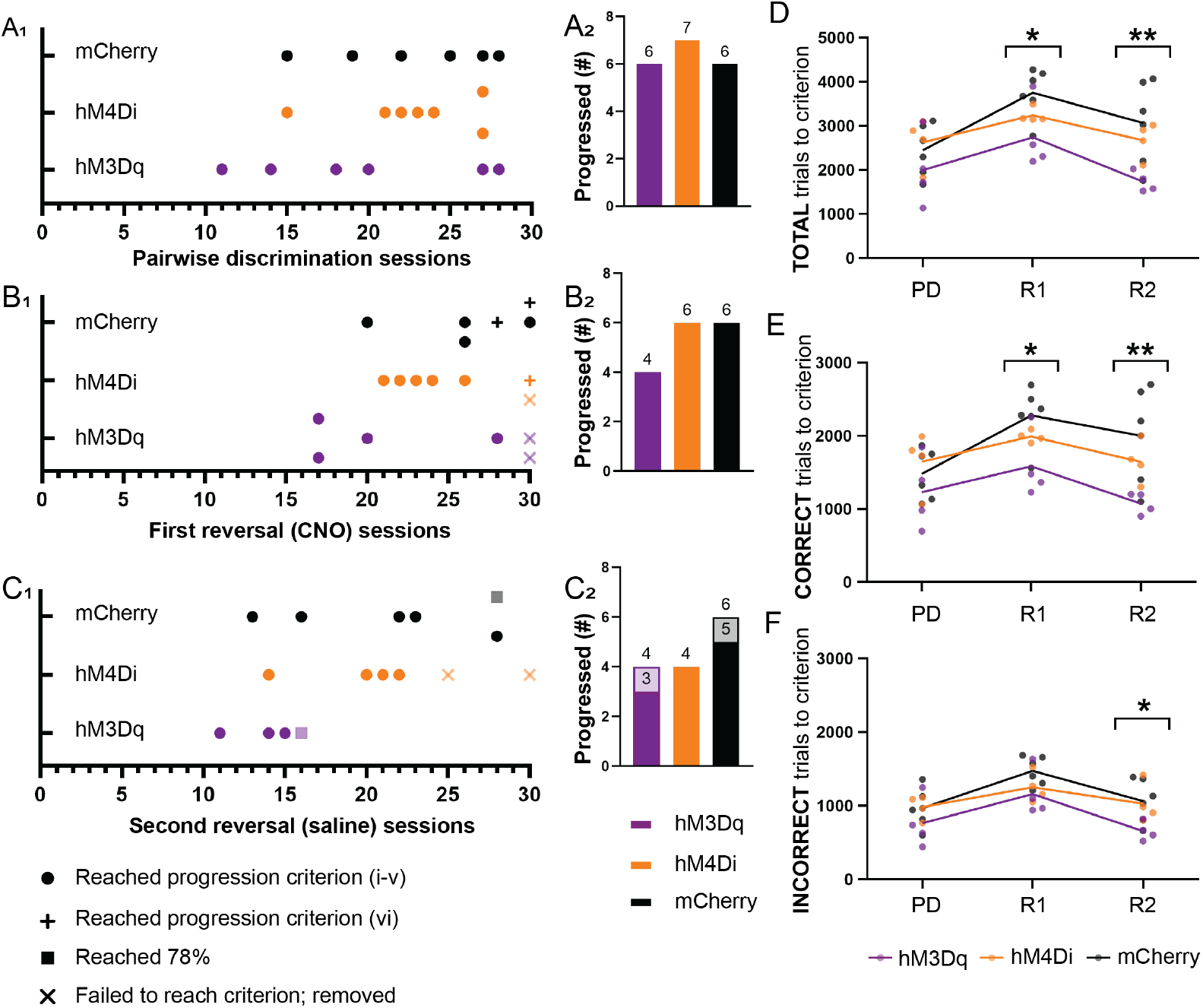
Duration of training required for each phase of visual discrimination and reversal learning. The number of sessions required (**1**) for rats to reach progression or removal criterion (**2**) in pairwise discrimination (PD; **A**), first reversal (R1; **B**) and second reversal (R2; **C**) for rats expressing either only mCherry (black), the activating hM3Dq-mCherry-DREADD (purple) or the inhibiting hM4Di-mCherry-DREADD (orange) in neuronal projections from the mPFC to the AcbSh. The total number of trials (**D**), as well as their distribution between correct (**E**) and incorrect (**F**) responses, required to reach criterion within each phase of reversal learning, for those rats that successfully completed all phases (hM3Dq n=4, hM4Di n=4, mCherry n=6). **D-F)** Two-way ANOVA followed by Tukey’s post-hoc multiple comparisons, see **Supplementary Table 3** for details. The total number of trials (**D**; *p*<.0001), number of correct trials (i.e. rewards collected; **E**; *p*=.0006), and number of incorrect trials (i.e. correction trials; **F**; *p*=.0003) required to reach criterion were all significantly different across phases. There was also a significant main effect of group for total trials (**D**; *p*=.0447 [hM3Dq < mCherry *p*=.0384]) and correct trials (**E**; *p*=.0424 [hM3Dq < mCherry *p*=.0364]). **D)** hM3Dq required significantly fewer total trials than mCherry in both R1 (*p*=.0468) and in R2 (*p*=.0067). **E)** hM3Dq required significantly fewer correct trials than mCherry in both R1 (*p*=.0350) and R2 (*p*=.0040). **F)** hM3Dq required significantly fewer incorrect trials in R2 than mCherry (*p*=.0446). Black vertical brackets show significant main effect group differences. Black horizontal brackets show significant between group comparisons within the enclosed phase (i.e. simple effects within phase). \**p*<.05, \*\**p*<.01.

## Discussion

The data presented here define a role for the mPFC-AcbSh pathway in mediating pathological body weight loss in the ABA model and suggest a neurobiological link between cognitive-behavioral flexibility and susceptibility to ABA. Specifically, chemogenetic suppression of the mPFC-AcbSh pathway allowed rats to maintain body weight during exposure to ABA conditions and prevented perseverative responding to changing reward contingencies in a touchscreen reversal learning paradigm. The maintenance of body weight was underpinned by an increase in food intake and food anticipatory activity (FAA) without a corresponding increase in overall running wheel activity. Importantly, modulation of the mPFC-AcbSh pathway in conditions of *ad libitum* feeding did not affect food intake, running or general locomotor activity, supporting the notion that these effects are specific to the expression of maladaptive behavior in ABA. These findings provide the first evidence that a neural circuit impacting both food intake *and* behavioral flexibility is involved in pathological weight loss in an animal model of AN.

The attenuation of body weight loss in ABA when mPFC-AcbSh pathway activity is suppressed aligns with multiple clinical studies in patients with AN that show excessive activity in regions of the prefrontal cortex. Prefrontal hyperactivity is demonstrated in AN patients compared to healthy weight controls in response to viewing negative emotional stimuli [43], in anticipation of monetary reward and feedback in an instrumental motivation task [9], as well as during food-cue association and reversal learning [44], supporting the view that both AN and ABA phenotypes involve a component of excessive cognitive control over emotional and reward processes. Similarly, the improvements in cognitive flexibility following suppression of prefrontal activity are in keeping with the previously established role of the rat prefrontal cortex in this behavioral phenotype [45, 46]. Together, these data suggest that flexible behavior *and* prefrontal activity are involved in the capacity, under ABA conditions, to maintain body weight. This series of studies highlights the potential contribution of a distinct fronto-striatal network to the excessive control over feeding behavior in patients with AN and further support the parallelism of the human condition and the rodent ABA model.

Although not addressed in this study, it is intriguing to consider the neurochemical basis of behavioral outcomes after modulation of prefrontal circuits. One of the obvious candidates is dopamine signaling; however, the published data in relation to this transmitter are not consistent. It is widely accepted that the majority of projections to and from the PFC are excitatory and glutamatergic [47] and that stimulation of PFC increases dopamine release in the nucleus accumbens (NAc) [48]. Increased striatal dopamine is associated with spontaneous locomotor hyperactivity [49], perhaps contributing to elevated ABA-associated running activity following excitation of the mPFC-AcbSh pathway. However, this same manipulation did not impact food intake during ABA or behavioral flexibility. The extensive colocalization between mCherry-expressing control neurons and Fos, described in the present study, suggests that a high level of baseline mPFC activity may create a “ceiling effect” for these measures, whereby further excitation of already high baseline activity produces fewer observable differences in behaviour than does suppression. Within the ABA paradigm, extracellular dopamine in the NAc is elevated during feeding [50] and chemogenetic stimulation of dopaminergic neurons in the ventral tegmental area projecting to the AcbSh increases food intake and FAA to prevent body weight loss [42]. This would support the notion that *increased* dopamine signaling in the accumbens is protective against ABA, however, both non-selective pharmacological *antagonism* of dopamine receptors [51] *and* selective dopamine receptor 2/3 (D2/3R) antagonism [52] also increases food intake in the ABA model. Similarly, the improvements in behavioral flexibility following mPFC-AcbSh inhibition are consistent with *decreased* dopamine 1 receptor (D1R) activity in the AcbSh, considering that pharmacological blockade of D1R in AcbSh selectively reduces early perseverative errors in this task [53]. The polarity of these findings makes it difficult to assign, with certainty, the role that dopamine plays in the interaction of prefrontal and accumbal circuitry. Perhaps this is unsurprising, in light of the large heterogeneity that exists in striatal dopamine neuron subpopulations and their connectivity. Dopamine release patterns are also variable (tonic, phasic or ramp-like) and do not necessarily align with dopamine neuron activity, suggesting local refinement of release characteristics within the striatum [54]. A more complete understanding of the role of dopamine is essential and may come from the application of approaches, including fiber photometry, where dopamine flux can be assessed in distinct neuronal populations during discrete behaviors.

Finally, impaired cognitive flexibility is beginning to be considered a transdiagnostic process that is common across eating disorders [55] and is associated with poorer improvement in social and occupational functioning in young people with emerging mental disorders [56]. The potential to treat specific cognitive and behavioral symptoms in patients with a range of psychiatric diagnoses (e.g. schizophrenia, substance use disorders, OCD, AN) is supported by the Research Domain Criteria (RDoC) concept to develop, for research purposes, new ways of classifying mental disorders based on dimensions of observable behavior and neurobiological measures [57, 58] and may afford progress in the understanding and treatment of mental disorders [59]. The discovery of a neurobiological link between cognitive flexibility and pathological body weight loss in an animal model of AN, as described here, highlights the utility of the ABA model for investigating the neurobiological bases of other cognitive and behavioral endophenotypes related to AN and how to treat them. Indeed, preliminary evidence from transcranial magnetic stimulation studies supports the notion that selective targeting of prefrontal cortical activity could aid in the treatment of AN, although effects may be transient [60] or predominantly associated with improvements in mood rather than core eating disorder symptoms [61]. More broadly, considerations of cognitive flexibility and ways to modulate it may have important implications across a much wider range of psychiatric conditions.

## Acknowledgements

These studies were supported by a Project Grant awarded by the Rebecca L Cooper Medical Research Foundation to CJF (PG2019373). We acknowledge the use of facilities at Monash Microimaging, Monash University, Australia and BioRender.com for images used in this manuscript. We also acknowledge Dr Emily Jaehne for technical assistance.

## Author contributions

CJF and BJO conceived of the study and wrote the manuscript. LKM, PNM, EG and CJF designed and performed the experiments and analyzed the data. LKM and PNM wrote parts of the manuscript. DCS and MVDB contributed to data interpretation and provided feedback on the manuscript. All authors approved the final version.

