## Supplementary Information for "Suppression of cortico-striatal circuit activity improves cognitive flexibility and prevents body weight loss in activity-based anorexia in rats"

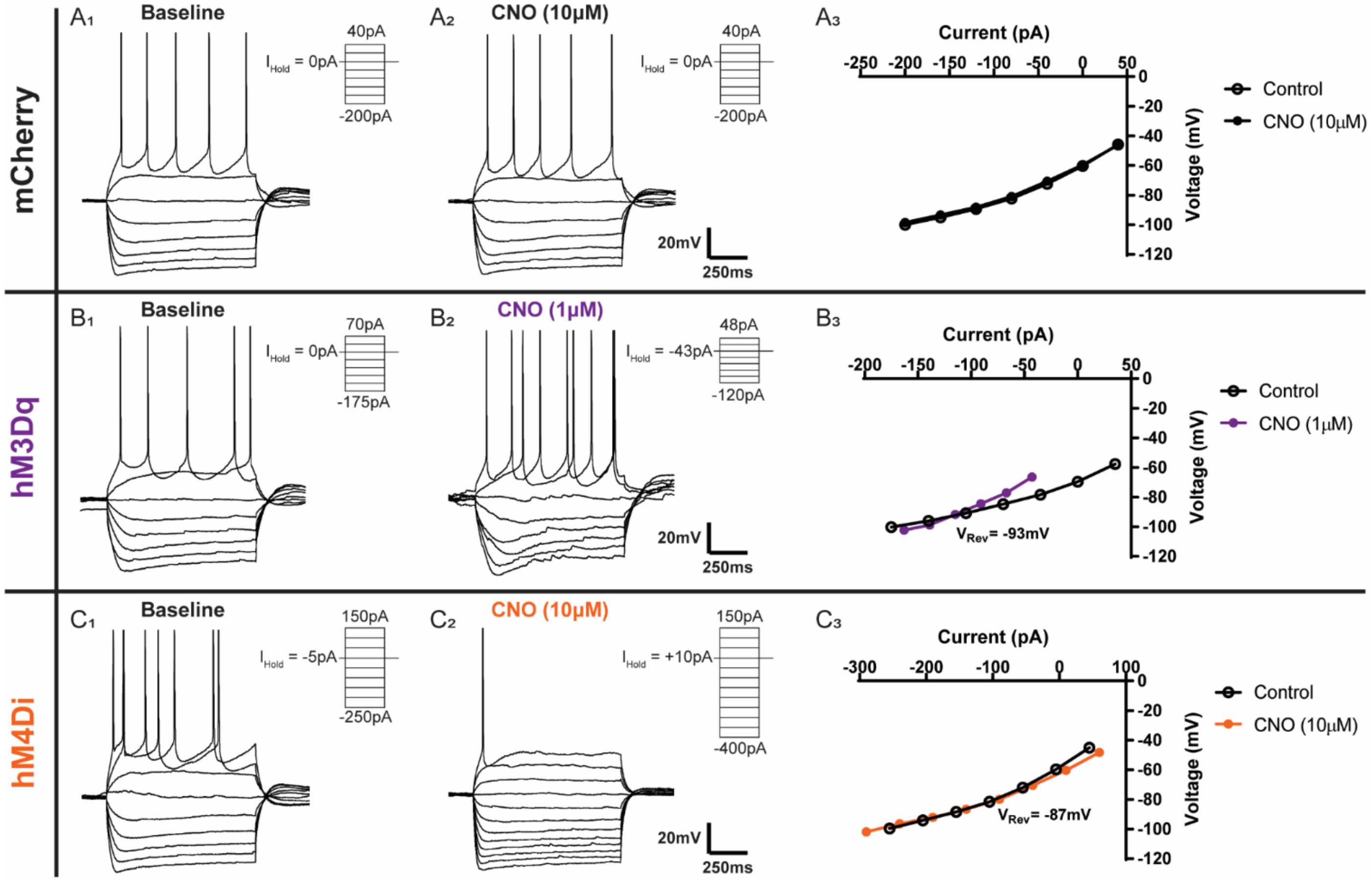


**Supplementary Figure 1: Current-voltage relationships for mCherry-, hM3Dq- and hM4Di-expressing mPFC-AcbSh neurons**

Whole-cell patch clamp electrophysiological recordings of **(A)** mCherry-expressing, **(B)** hM3Dq-expressing and **(C)** hM4Di-expressing mPFC to AcbSh pyramidal neurons. A series of current steps were injected (as indicated, inset) into cells at **(1)** baseline and **(2)** at the peak of the response to bath-applied CNO (1-10μM). Current-voltage relationship plots were constructed, revealing for hM3Dq-expressing cells **($B_3$)** a CNO-induced increase in input resistance with a reversal potential of ≈-93mV, and for hM4Di-expressing cells **($C_3$)** a CNO-induced reduction in input resistance with a reversal potential of ≈-87mV, indicating the closure and opening of one or more potassium channels, respectively.

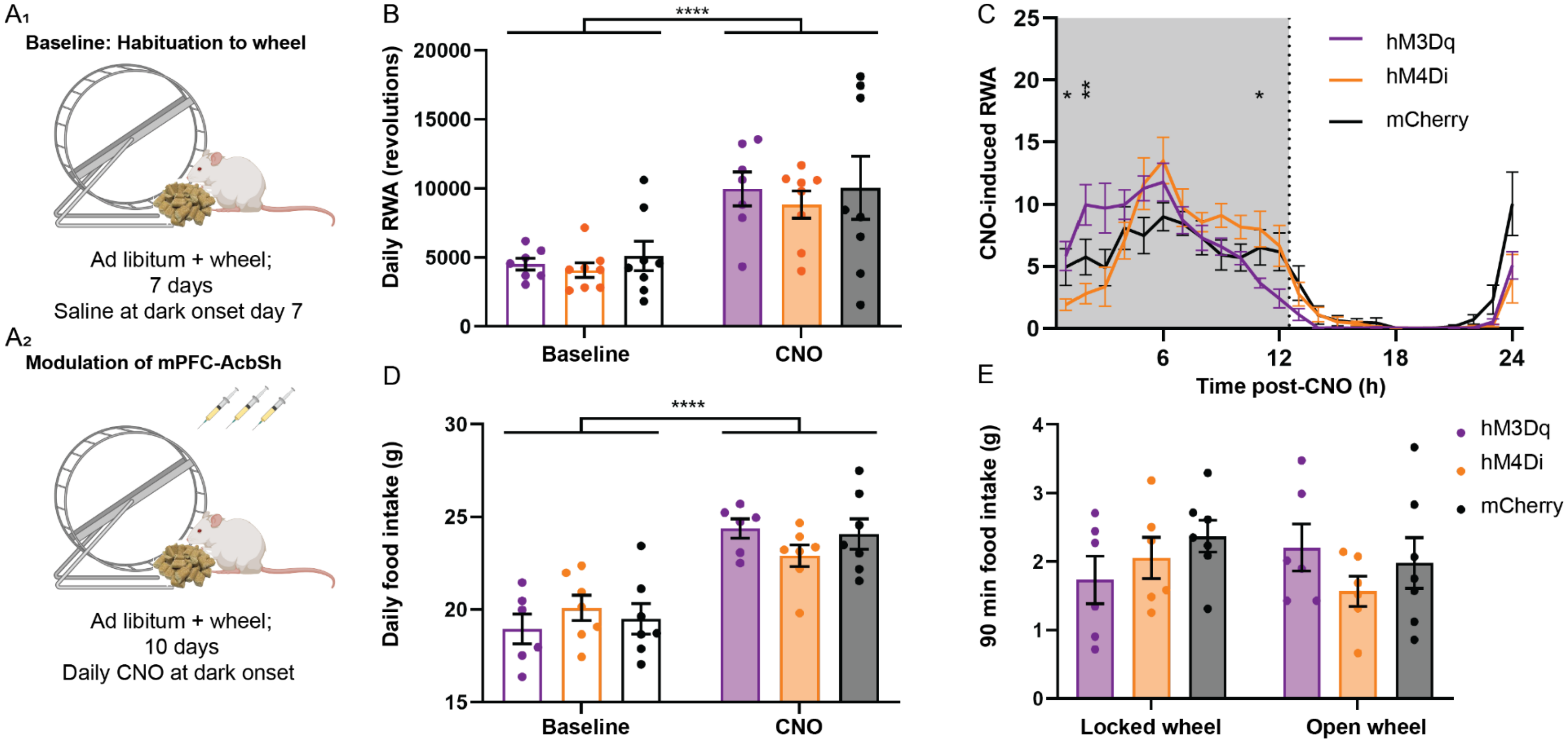


**Supplementary Figure 2: Effects of mPFC-AcbSh modulation on activity and food intake with *ad libitum* food access**

Outcomes for rats maintained on ad libitum food access and exposed to wheel access and CNO administration that mirrors the ABA protocol **(A)** expressing either only mCherry (black, n=7-9), activating hM3Dq-mCherry DREADDs (purple, n=6-7) or inhibiting hM4Di-mCherry DREADDs (orange, n=7-9) in neurons projecting from the mPFC to the AcbSh. Daily RWA **(B)** and daily food intake **(D)** significantly increased from baseline to CNO for the entire cohort (both *p*s<.0001) and each individual group (all *p*s≤.0016), indicating an effect of time rather than CNO, evidenced by the mCherry group and in keeping with the gradual increase in both measures for all groups across days shown in **Figure 3 B and C. C)** Over the 10-day CNO phase mean hourly RWA (% of daily RWA) was significantly different over the 24 x 1 hour blocks with a significant time*group interaction (both *p's*<.0001), with hM3Dq running significantly more than hM4Di at 1h (*p*=.0352) and 2h (*p*=.0089) post-CNO, and hM3Dq running significantly less than hM4Di at 11h post-CNO (*p*=.0461). **E)** There were no significant main effects or interaction on 90-minute food intake following CNO administration (all *p*s≥.1652). **B-E)** Two-way ANOVA followed by post-hoc Tukey's multiple comparisons (see **Supplementary Table 3** for details); individual animals (dots) with group mean (bars/lines) ± SEM. *$p$<.05, **$p$<.01, ****$p$<.0001.

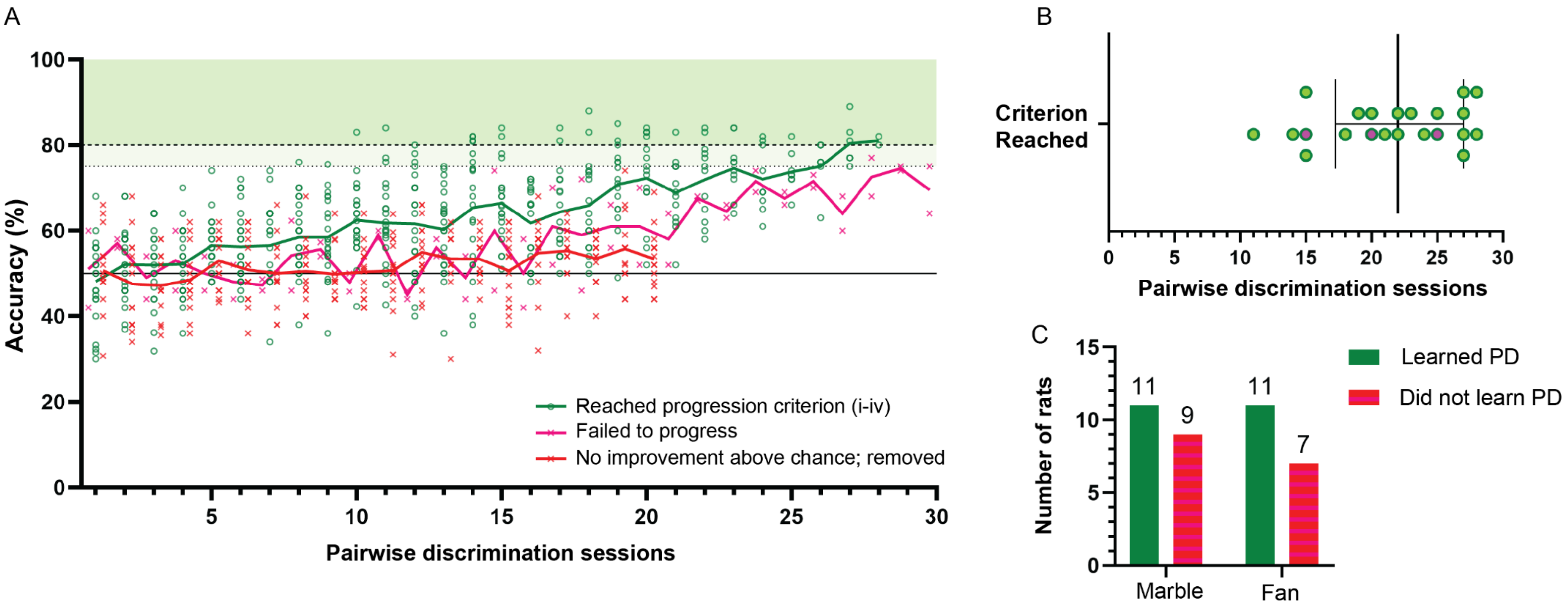


**Supplementary Figure 3: Visual discrimination learning rates**

**A)** Accuracy (% correct in accuracy trials [AT]) in Pairwise Discrimination (PD) for rats that i) reached progression criterion i-iv (see Supplementary Table 2 for details; green lines and circles; n=22), ii) were removed after 20 sessions as performance remained no better than chance (red lines and crosses; n=14), and iii) showed evidence of learning (performing better than chance after 20 sessions) but failed to reach progression criterion after the maximum of 30 sessions (failed to progress, pink lines and crosses; n=2). **B)** Pairwise discrimination sessions (one per day, 6 days a week) required for 22 rats to achieve progression criterion (Median (long vertical line; 22.00) with IQR (17.25 to 27). Of these 22 rats, 3 had off target/no DREADDs (bright pink filled circles) and were excluded from analysis, leaving n=19 with on target DREADDs (bright green filled circles). **C**) Achievement of PD progression criterion was not influenced by which image was rewarded, Fisher's exact test (two-tailed) *p*=.7521.

**Supplementary Methods 1: Brain slice preparation for electrophysiology**

Three weeks after viral injections, rats were anaesthetised (5% isoflurane) and decapitated. Brains were rapidly collected into ice-cold sucrose-modified artificial cerebrospinal fluid (in mM, 200 Sucrose, 1.9 KCl, 1.2 $NaH_2PO_4$, 26 $NaHCO_3$, 5.0 D-glucose, 5.0 D-mannitol, 2.0 sodium pyruvate, 0.5 ascorbic acid, 10 $MgCl_2$), and were cut into 300µM coronal sections of the mPFC using a vibratome (VT1000S; Leica, Cambridge, UK). Slices were transferred to recording aCSF (in mM, 127 NaCl, 1.9 KCl, 1.2 $KH_2PO_4$, 26 $NaHCO_3$, 5.0 D-glucose, 5.0 D-mannitol, 1.3 $MgCl_2$, 2.4 $CaCl_2$, 0.34 ascorbic acid; equilibrated with 95% O2 and 5% CO2, pH 7.3-7.4, 300-305mOsm/L) at 34-35°C for 25 mins, and then left to rest at room temperature for 60 mins before electrophysiological recordings were conducted.

**Supplementary Methods 2: Whole-cell patch clamp validation of DREADD function**

For electrophysiological recordings, mPFC brain slices were transferred to a recording chamber and constantly bathed in aCSF (flow rate = 3-6mL/min). DREADD(mCherry)-expressing neurons were identified using epifluorescence, then with differential interference contrast microscopy healthy neurons were selected for recording using a Multiclamp 700A amplifier (Axon Instruments, Foster City, CA, USA). Patch-pipettes were pulled using a horizontal puller (Sutter Instruments, USA; P1000 model) from thin-walled borosilicate glass (Harvard Apparatus; GC150-TF10) to resistances between 5 and 10MΩ when filled with intracellular recording solution (in mM, 140 K-gluconate, 10 HEPES, 10 KCl, 1.0 EGTA and 2.0 $Na_2ATP$, 0.3 $Na_2GTP$; pH adjusted to 7.35 with KOH, osmolarity adjusted to ~305mOsm/L with sucrose). In current clamp mode, baseline recordings were taken until membrane potential was stable (~5 minutes), then CNO dissolved in recording aCSF from a stock solution was bath-applied to the brain slice (50mM stock solution in DMSO; final concentration 1µM for hM3Dq-expressing neurons or 10µM for mCherry- and hM4Di-expressing cells, approximating relative *in vivo* doses). The time to peak response following CNO application was typically 3-5 minutes, after which current injections were made to determine current-voltage relationships, input resistance and rheobase. Membrane potential was recorded when stable just prior the application of CNO, and at the peak of the CNO response. Firing frequency was determined by measuring the number of action potentials fired in the 60 seconds following the peak CNO response (hM3Dq-expressing cells). Input resistance was determined by the membrane voltage response to a small hyperpolarising step (-30 to -60pA) while cells were held at approximately -60mV. Rheobase was obtained by holding cells to -60mV, then injecting positive current steps of 2pA increment until the cell first fired at two concordant current measures. For data analysis, the signal was digitised at 2-10 kHz and analysed utilising pClamp10 (Axon Instruments).

**Supplementary Methods 3: Touchscreen Training**

Rats were trained once per day 6 days per week during the dark phase. Rats were placed in a trapezoid operant conditioning chamber (Campden Instruments; 25cm wide at touchscreen, 13cm wide at magazine, 33cm long between screen and magazine, 30cm high) with the touchscreen (15in diagonal, rotated) covered by a black plastic mask with two 10 x 10 cm openings (bottom edge of opening is 16 cm from floor) through to the touchscreen that differentiates the image locations from the background and helps prevent accidental screen touches. Rewards consisted of 45mg sucrose pellets delivered individually via a pellet dispenser into the magazine (body housed outside the chamber, opening in the wall opposite the touchscreen), which has its own light and IR beam to detect entry of the rat's nose. There is also a house light, tone generator and click generator, all of which act as conditioned reinforcers as described below. All data were acquired using ABET II Touch software (Lafayette Instruments; IN, USA).

There are multiple pre-training steps (maximum 60 minutes per session) that progressively teach the rats the steps involved in the task (for details see [1] and **Supplementary Table 1**). Briefly, rats 1) are habituated to the sucrose pellets (~20 per rat given in home cage) and touchscreen chambers (Habituation), 2) learn to associate the touchscreen images with sucrose rewards (Initial Touch), 3) learn to touch the image to receive a reward (Must Touch), 4) learn to initiate the presentation of the image/s (Must Initiate), and 5) learn the 'punishment' for making an incorrect response (Punish Incorrect). Each rat received 2 i.p. injections of saline 30 minutes before the punish incorrect session on 2 consecutive days in their final week of pretraining for habituation. Following successful completion of pretraining rats were progressed to the task of interest, reversal learning.

Our reversal learning task consisted of 3 phases each with a maximum 30 sessions. Each session (max 60 minutes) required rats to correctly respond to 50 or 100 trials (within each phase, pairs of 50 trial sessions were initially given until rats performed quickly enough to complete 100 trials within 60 minutes): a trial consisted of repeated presentations of the two images (pinwheel fan and marble array, see **Figure 3A**), one of which was pseudo randomly designated as correct (balanced across treatment groups and number of pretraining sessions), in the same location (left/right) until the correct image was touched which was rewarded with delivery of a sucrose pellet and initiated progression to the next (accuracy) trial. Only the first presentation in each trial (accuracy trial [AT]) contributed towards the accuracy calculation (% correct); subsequent identical image presentations following an incorrect touch are termed correction trials (CT) and do not contribute towards accuracy but served as a learning reinforcer and are repeated until the correct image is touched. During the initial Pairwise discrimination (PD) phase rats progressively learned to identify and touch the correct image. Over the last ~week of PD sessions rats received i.p. injections of saline 30 minutes before the start of their session. The reward contingency of the images was then switched (First reversal [R1] phase) and daily CNO (0.3-3.0mg/kg) administration was initiated via i.p. injection 30 minutes before each testing session throughout this phase. A serial reversal (Second reversal [R2] phase) followed in which the reward contingency reverted to that used in PD; CNO administration ceased and was replaced by i.p. saline. Each phase continued until rats reached progression criterion (see **Supplementary Table 2** for details), or removal criterion (performance no better than chance after 20 sessions, or failing to reach progression criterion after the maximum allowed 30 sessions).

**Supplementary Table 1: Details of training steps used in touchscreen operant training**

| Stage | Image/s | Response | Tone | Magazine light | Sucrose pellet/s | House light | Magazine click | Next trial? | Trials completed count | Progression criterion |
|---|---|---|---|---|---|---|---|---|---|---|
| **Habituation** | N/A | N/A | N/A | N/A | ~20 in magazine before session | Off | N/A | N/A | N/A | Consume all pellets in 60 minutes on two consecutive sessions |
| **Initial Touch** | One image, one blank window<br><br>Image displayed until touched or for max of 30 seconds | Touch image | Yes –when image touched | Turns on with pellet delivery – turns off when reward collected | 3, collection starts 20s ITI | Off | N/A | Normal – Automatic start after 20s ITI | +1 | Complete 100 trials in 60 minutes once or max 3 sessions |
| | | Image not touched after 30s | Yes –when image disappears after 30s | | 1, collection starts 20s ITI | Off | N/A | | +1 | |
| | | Touch to blank window during trial | No effect | | | | | | | |
| **Must Touch** | One image, one blank window<br><br>Image displayed until touched | Touch image | Yes –when image touched | Yes – turns off when reward collected | 1, collection starts 20s ITI | Off | N/A | Normal – Automatic start after 20s ITI | +1 | Complete 100 trials in 60 minutes once |
| | | Touch to blank window during trial | No effect | | | | | | | |
| **Must Initiate** | One image, one blank window | Image touch | Yes –when image touched | Yes – turns off when | 1, collection | Off | Yes – with trial initiation | Normal - Must be initiated | +1 | Complete 100 trials in 60 minutes |

| | | | | | | | | | | |
|---|---|---|---|---|---|---|---|---|---|---|
| | Image displayed until touched | | | reward collected | starts 20s ITI | | | with magazine nose poke after 20s ITI | | once in first 3 sessions; Complete >90 trials in 60 minutes in 4th session |
| | | Blank touch | No effect | | | | | | | |
| **Punish Incorrect** | One image, one blank window<br><br>Image displayed until image touch or blank touch | Touch image | Yes –when image touched | Yes – turns off when reward collected | 1, collection starts 15s ITI | Off | Yes – with trial initiation | Accuracy trial - Must be initiated after ITI | +1 | Complete 100 trials in 60 minutes, with accuracy ≥80%for accuracy trials in two sessions |
| | | Blank touch | No | No | 0 | On for 5s time out; turns off after timeout and 15s ITI begins | Yes – with trial initiation following 5s time-out and 15s ITI | Correction trial - Must be initiated after ITI | +0 | |
| **Pairwise discrimination + reversals** | Fan/pinwheel in one window and marble array in other window<br><br>pseudorandom side allocation* | Correct image touched | Yes –when image touched | Yes – turns off when reward collected | 1, collection starts 15s ITI | Off | Yes – with trial initiation | Accuracy trial - Must be initiated after ITI | +1 | **See Supp Table 2 below** |
| | | Incorrect image touched | No | No | 0 | On – 5s time out; turns off after timeout and 10s ITI begins | Yes – with trial initiation following 5s time-out and 10s ITI | Correction trial - Must be initiated after ITI | +0 | |

* Image will not be displayed on the same side more than 3 consecutive trials (excluding correction trials, where image+side is repeated until the correct response is made)

**NOTE**: an incorrect response does not add to the count of completed trials. **NOTE**: ONLY the response to the first image presentation within each trial (accuracy trial, AT) contributes towards the accuracy calculation, i.e. correct responses to correction trials are rewarded to facilitate learning but do not contribute towards accuracy. **NOTE**: Initially rats are very slow to complete the required number of trials in punish incorrect and all three phases of the reversal learning task as they require a lot of correction trials. Therefore, pairs of 50 trial sessions are given until rats are quick enough to complete the required 100 trials in 60 minutes. Additionally, to facilitate the completion of the required number of trials we shortened the inter-trial interval (ITI) from 20s to 15s following a correct response and following an incorrect response to 15s in punish incorrect and 10s in the reversal learning task.

**Supplementary Table 2: Progression criteria for touchscreen cognitive testing**

| Phase | Sessions | Progression criteria. The session in which a rat achieves any one of: |
|---|---|---|
| **Pairwise Discrimination** | 1-30 | **i)** second consecutive session with accuracy ≥ 80%<br>**ii)** accuracy ≥75% in session immediately following one with accuracy ≥80%<br>**iii)** a second session with accuracy ≥80% following accuracy dropping below 75% after previous session with accuracy ≥80% |
| | 26-30 | **i-iii)**<br>**iv)** accuracy ≥80% in session immediately following one with accuracy ≥75% |
| **First reversal** | 1-20 | **i-iii)** |
| | 21-25 | **i-iv)** |
| | 26-30 | **i-iv)**<br>**v)** accuracy ≥80%<br>**vi)** accuracy ≥78% |
| **Second reversal** | 1-20 | **i-iii)** |
| | 21-30 | **i-iv)** |

**Supplementary Table 3: Statistical analysis details**

| Figure | Statistics test details |
|---|---|
| **2D** | One-way ANOVA: F(2, 32) = 4.137, ***p*=.0252**<br>• hM3Dq v mCherry *p*=.7756; hM3Dq < hM4Di ***p*=.0229;** hM4Di v mCherry *p*=.1036 |
| **2F** | One-way ANOVA: F(2, 32) = 6.063, ***p*=.0059**<br>• hM3Dq > mCherry ***p*=.0126**; hM3Dq > hM4Di ***p*=.0190**; hM4Di v mCherry *p*>.9999 |
| **2G** | One-way ANOVA: F(2, 32) = 3.779, *p*=.0336<br>• hM3Dq v mCherry *p*>.9999; hM3Dq v hM4Di *p*=.1284; hM4Di < mCherry ***p*=.0383** |
| **4D** | Mixed effects analysis – Geisser-Greenhouse's epsilon .7110<br>Main effect of group: F(2, 13) = 3.834, ***p*=.0491**<br>• R1: hM4Di > mCherry ***p*=.0366**, hM4Di > hM3Dq ***p*=.0121**, hM3Dq v mCherry *p*=.8043<br>Main effect of phase: F(1.422, 18.49) = 53.40***, p*<.0001**; PD > R1 ***p*<.0001**, PD > R2 ***p*<.0001**, R1 < R2 ***p*=.0161**<br>• hM3dq: PD > R1 ***p*=.0096**, R1 < R2 ***p*=.0346**<br>• hM4Di: PD > R1 ***p*=.0433**, PD > R2 ***p*=.0095**<br>• mCherry: PD > R1 ***p*=.0065**, R1 < R2 ***p*=.0304**<br>Group*phase interaction: F(4, 26) = 2.586, *p*=.0604 |
| **4E** | Mixed effects analysis – Geisser-Greenhouse's epsilon .8571<br>Main effect of group: F(2, 13) = 1.715, *p*=.2183<br>• R1: hM4Di < hM3Dq ***p*=.0025**, hM3Dq > mCherry ***p*=.0346**<br>Main effect of phase: F(1.714, 22.29) = 19.73***, p*<.0001**; R1 > R2 ***p*=.0013**<br>• hM3Dq: PD < R1 ***p*=.0254**, R1 > R2 ***p*=.0054**<br>• mCherry: R1 > R2 ***p*=.0106**<br>Group*phase interaction: F(4, 26) = 4.828, ***p*=.0048** |
| **4F** | Mixed effects analysis – Geisser-Greenhouse's epsilon .8854<br>Main effect of group: F(2, 13) = 3.117, *p*=.0784<br>• R1: hM4Di < hM3Dq ***p*=.0008**, hM3Dq > mCherry ***p*=.0382**<br>Main effect of phase: F(1.771, 23.02) = 49.66, ***p*<0.0001;** PD < R1 ***p*=.0004**, R1 > R2 ***p*=.0015**<br>• hM3Dq: PD < R1 ***p*=.0108**, R1 > R2 ***p*=.0032**<br>• mCherry: PD < R1 ***p*=.0089**, R1 > R2 ***p*=.0086**.<br>Group*phase interaction: F(4, 26) = 9.196, ***p*<0.0001** |
| **4G** | Mixed effects analysis – Geisser-Greenhouse's epsilon .9780<br>Main effect of group: F(2, 13) = 0.05693, *p*=.9449<br>Main effect of phase: F(1.956, 25.43) = 30.43, ***p*<0.0001**; PD < R1 ***p*<.0001**, PD < R2 ***p*=.0045**, R1 > R2 ***p*=.0167** |

| | |
|---|---|
| | • hM3Dq: PD < R1 ***p*=.0339**, R1 > R2 ***p*=.0004**<br>• hM4Di: PD > R2 ***p*=.0465**<br>• mCherry PD < R1 ***p*=.0009**, R1 > R2 ***p*=.0211**<br>Group*phase interaction: $F(4, 26) = 2.683$, *p*=.0537 |
| **4H** | Mixed effects analysis (*incorrect* latency) – *sphericity assumed*, Geisser-Greenhouse's epsilon .9858<br>Main effect of group: $F(2, 13) = 1.289$, *p*=.3085<br>• R1: hM3Dq > hM4Di ***p*=.0148**, hM3Dq > mCherry ***p*=.0193**<br>Main effect of phase: $F(2, 26) = 1.068$, *p*=.3583<br>• hM3Dq: PD < R1 ***p*=.0040**<br>• mCherry: PD > R2 ***p*=.0443**<br>Group*phase interaction: $F(4, 26) = 4.770$**, *p*=.0051**<br><br>Mixed effects analysis (*correct* latency) – *sphericity assumed*, Geisser-Greenhouse's epsilon .8305<br>Main effect of group: $F(2, 13) = 2.835$, *p*=.0951<br>• R1: hM3Dq > hM4Di ***p*=.0018**, hM3Dq > mCherry ***p*=.0014**<br>Main effect of phase: $F(2, 26) = 3.641$, ***p*=.0404**; R1 > R2 ***p*=.0317**<br>• hM3Dq: PD < R1 ***p*=.0013**, R1 > R2 ***p*=.0235**<br>• mCherry: PD > R2 ***p*=.0248**<br>Group*phase interaction: $F(4, 26) = 5.161$, ***p*=.0034** |
| **4I** | Mixed effects analysis) – *sphericity assumed*, Geisser-Greenhouse's epsilon .9977<br>Main effect of group: $F(2, 13) = 4.034$, ***p*=.0434**; hM3Dq > hM4Di ***p*=.0404**<br>• R2: hM3Dq > hM4Di ***p*=.0365**<br>Main effect of phase: $F(2, 26) = 14.19$, ***p*<0.0001**; PD > R1 ***p*=.0004**, PD > R2 ***p*=.0002**<br>• hM4Di: PD > R2 ***p*=.0270**<br>• mCherry: PD > R1 ***p*=.0072**, PD > R2 ***p*=.0030**<br>Group*phase interaction: $F(4, 26) = 0.2722$, ***p*=.8932** |
| **5D** | Mixed effects analysis) – *sphericity assumed*, Geisser-Greenhouse's epsilon .8253<br>Main effect of group: $F(2, 11) = 4.176$, ***p*=.0447**; hM3Dq < mCherry ***p*=.0384**<br>• R1: hM3Dq < mCherry ***p*=.0468**<br>• R2: hM3Dq < mCherry ***p*=.0067**<br>Main effect of phase: $F(2, 22) = 14.69$, ***p*<0.0001**; PD < R1 ***p*=.0001**, R1 > R2 ***p*=.0009**<br>• hM3Dq: R1 > R2 ***p*=.0135**<br>• mCherry: PD < R1 ***p*=.0002**, R1 < R2 ***p*=.0424** |

| | |
|---|---|
| | Group*phase interaction: F(4, 22) = 1.498, *p*=.2372 |
| **5E** | Mixed effects analysis) – *sphericity assumed*, Geisser-Greenhouse’s epsilon .8953<br>Main effect of group: F(2, 11) = 4.271, ***p*=.0424**; hM3Dq < mCherry ***p*=.0364**<br>• R1: hM3Dq < mCherry ***p*=.0350**<br>• R2: hM3Dq < mCherry ***p*=.0040**<br>Main effect of phase: F(2, 22) = 10.49, ***p*=.0006**; PD < R1 ***p*=.0006**, R1 > R2 ***p*=.0085**<br>• mCherry: PD<R1 ***p*=.0003**, PD < R2 ***p*=.0153**<br>Group*phase interaction: F(4, 22) = 2.006, *p*=.1289 |
| **5F** | Mixed effects analysis – Geisser-Greenhouse’s epsilon .7384<br>Main effect of group: F(2, 11) = 3.142, *p*=.0833<br>• R2: hM3Dq < mCherry ***p*=.0446**<br>Main effect of phase: F(1.477, 16.24) = 16.87, ***p*=.0003**; PD < R1 ***p*=.0003**, R1 > R2 ***p*<.0001**<br>• hM3Dq: R1>R2 ***p*=.0497**<br>• mCherry: PD<R1 ***p*=.0259**, R1>R2 ***p*=.0041**<br>Group*phase interaction: F(4, 22) = 0.8521, *p*=.5077 |
| **S2B** | Two-way ANOVA<br>Main effect of group: F(2, 20) = 0.2711, *p*=.7653<br>Main effect of phase: F(1, 20) = 57.41, ***p*<.0001**<br>• hM3Dq: ***p*=.0006**<br>• hM4Di: ***p*=.0013**<br>• mCherry: ***p*=.0009**<br>Group*phase interaction: F(2, 20) = 0.09735, *p*=.9077 |
| **S2C** | Two-Way ANOVA – Geisser-Greenhouse’s epsilon =.1922<br>Main effect of group: F(2, 20) = 0.02558, *p*=0.9748<br>• 1h: hM3Dq > hM4Di ***p*=.0352**<br>• 2h: hM3Dq > hM4Di ***p*=.0089**<br>• 11h: hM3Dq < hM4Di ***p*=.0461**<br>Main effect of time: F(4.421, 88.43) = 35.31, ***p*<.0001**<br>Group*phase interaction:F(46, 460) = 2.545, ***p*<.0001** |
| **S2D** | Two-way ANOVA<br>Main effect of group: F(2, 17) = 0.05601, *p*=.9457<br>Main effect of phase: F(1, 17) = 118.3, ***p*<.0001**<br>• hM3Dq: ***p*<.0001**<br>• hM4Di: ***p*=.0016** |

| | |
|---|---|
| | • mCherry: ***p*<.0001**<br>Group*phase interaction: F(2, 17) = 3.790, ***p*=.0435** |
| **S2E** | Two-way ANOVA<br>Main effect of group: F(2, 16) = 0.5541, *p*=.5852<br>Main effect of wheel: F(1, 16) = 0.4060, *p*=.5330<br>Group*phase interaction: F(2, 16) = 2.019, *p*=.1652 |
